# Cell-ACDC: a user-friendly toolset embedding state-of-the-art neural networks for segmentation, tracking and cell cycle annotations of live-cell imaging data

**DOI:** 10.1101/2021.09.28.462199

**Authors:** Francesco Padovani, Benedikt Mairhörmann, Pascal Falter-Braun, Jette Lengefeld, Kurt M. Schmoller

## Abstract

Live-cell imaging is a powerful tool to study dynamic cellular processes on the level of single cells with quantitative detail. Microfluidics enables parallel high-throughput imaging, creating a downstream bottleneck at the stage of data analysis. Recent progress on deep learning image analysis dramatically improved cell segmentation and tracking. Nevertheless, manual data validation and correction is typically still required and broadly used tools spanning the complete range of live-cell imaging analysis, from cell segmentation to pedigree analysis and signal quantification, are still needed. Here, we present Cell-ACDC, a user-friendly graphical user-interface (GUI)-based framework written in Python, for segmentation, tracking and cell cycle annotation. We included two state-of-the-art and high-accuracy deep learning models for single-cell segmentation of yeast and mammalian cells implemented in the most used deep learning frameworks TensorFlow and PyTorch. Additionally, we developed and implemented a cell tracking method and embedded it into an intuitive, semi-automated workflow for label-free cell cycle annotation of single cells. The open-source and modularized nature of Cell-ACDC will enable simple and fast integration of new deep learning-based and traditional methods for cell segmentation or downstream image analysis.

**Source code:** https://github.com/SchmollerLab/Cell_ACDC

## Introduction

Live-cell imaging is a powerful technique that allows studying complex cellular dynamics by providing spatiotemporal information of subcellular events. Microfluidic devices that maintain constant environments enable parallel imaging of thousands of cells for many hours in a single experiment. However, downstream analysis typically involves many potentially time-consuming steps, e.g., cell segmentation, tracking, and pedigree annotation. Thus, for the large amount of data typically produced by a live-cell imaging experiment downstream extraction of biologically relevant information becomes the rate-limiting step.

While traditional segmentation algorithms had low generalization power, recent advances in deep learning, and specifically in fully convolutional neural networks (FCNN) based on U-Net^1^, have greatly enhanced segmentation accuracy and degree of automation^2,3^. More specifically, in the case of live-cell microscopy of yeast and other organisms (e.g., stem mammalian cells), two neural networks recently published (YeaZ^4^ and Cellpose^5^) drastically improve the segmentation process. However, even these neural networks do not achieve perfect segmentation, and – depending on the question – manual verification or correction of segmentation and tracking is often still essential for high-quality microscopy image analysis.

In addition, in most cases analysis of movies that image cells over multiple generations also requires the correct annotation of pedigrees and cell cycle transitions. This is particularly true for budding yeast, because the bud, even though still a part of the mother cell, is recognized as an individual object by most segmentation routines. To obtain information about a full cell cycle, it is therefore key to link a bud to its mother cell and determine the time point of cell division. Importantly, for budding yeast such cell cycle annotations can in part be performed in a label-free manner based on the phase contrast signal: bud emergence is linked to S-phase entry, and cell division is typically detectable by a sudden movement of the bud that is not mechanically linked to the mother cell anymore. Unfortunately, such pedigree and cell cycle annotation is still a completely manual process that requires careful inspection of every single frame to catch and annotate the time point of cell division. Fluorescent tagging of proteins that locate to the bud neck connecting mother and bud, or of histones to monitor S-phase and observe nuclear localization facilitates pedigree annotation and has been used for automation^6–9^. Nevertheless, even with endogenous tagging (which comes with the cost of requiring genetic manipulation as well as one fluorescent channel that otherwise could be used for other purposes), automated approaches do not achieve the accuracy required for many questions and thus still require manual inspection and correction.

Although many software tools dedicated to the analysis of budding yeast live-cell microscopy have been developed in the past (ImageJ/Fiji^10^, MorphoLibJ^11^, PhyloCell^12^, CellProfiler^13^, Cell Tracer^14^, Wood et al.^15,16^, Cell Star^17^, Cell Serpent^18^, CellID^19^, Tracker^20^, DISCO^21^, YeastSpotter^22^, YeastNet^23^, DeepCell^24^, Cellbow^25^), to the best of our knowledge, none of them spanned the entire image analysis pipeline from CNN-base segmentation to cell cycle analysis, and fluorescent signal quantification.

Here we present an open-source graphical user-interface (GUI)-based framework (written in Python 3) embedding state-of-the-art neural networks (YeaZ and Cellpose) and smart algorithms that allow for fast, replicable, and accurate microscopy image analysis. The tools provided cover the entire image analysis pipeline from the raw microscopy file to the final quantification of the feature of interest. We named this software Cell-ACDC for Cell-Analysis of the Cell Division Cycle. The pipeline was developed with a community-centred approach, where users from several research groups provided feedback and suggestions that were implemented into the pipeline. Cell-ACDC provides for the first time the possibility to constantly visualize and correct any segmentation, tracking, or cell cycle annotation error in a fast and intuitive way. It includes several smart algorithms and shortcuts that automatically propagate any change to past and future frames to continuously maintain data integrity and correctness. While designed for budding yeast, many of the included tools are ready to use for other model systems, such as human stem cells and fission yeast. If adopted by the community, Cell-ACDC could standardize handling and analysis of live-cell microscopy data and facilitate exchanging data between different labs.

## Results

Cell-ACDC provides a framework that allows the implementation of the entire image analysis pipeline, from the raw microscopy file to visualizing and plotting the results (Figure 1A). In the first steps, the raw microscopy files are converted into TIF files (one for each channel of each position) using the popular Bio-Formats^26^ library in a fully automated Python routine. Using Bio-Formats allows for standardized reading of the file metadata, such as the number of frames, the number of z-slices in a z-stack or the time interval between each frame etc. Furthermore, we provide full support for 2D, 3D (single z-stacks or 2D images over time) and 4D images (3D z-stacks over time) with multiple channels and multiple positions.

**Fig 1.**
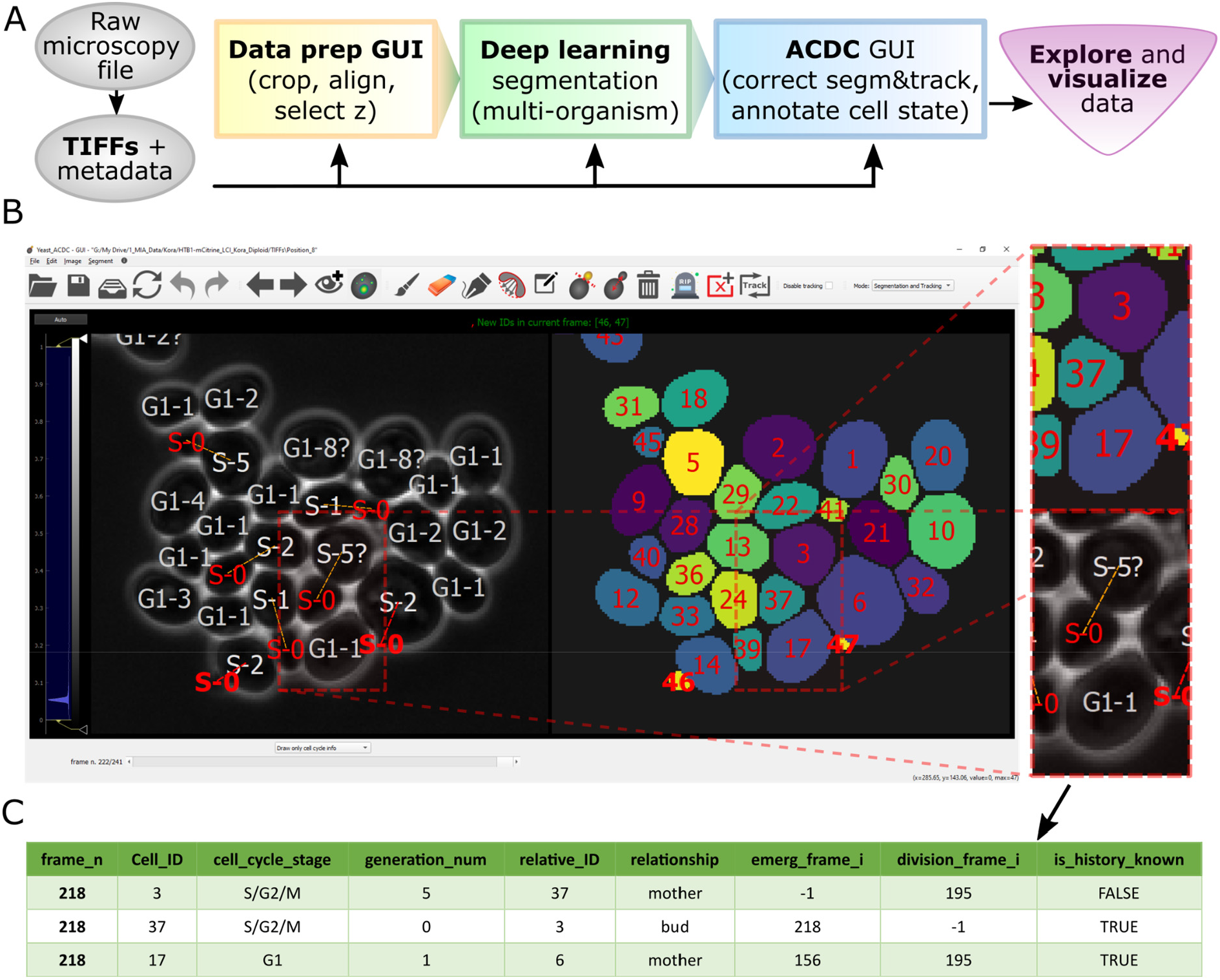
Overview of pipeline and GUI. (A) Flowchart representation of the Cell-ACDC pipeline. In the first step (either in Python or Fiji macro) the raw microscopy file(s) is/are automatically converted into TIFs, the relevant metadata is extracted, and the files are arranged in the data structure required by Cell-ACDC. Next, the user can launch any of the three main modules: 1) GUI-based data prep where the user can align time-lapse data, select a z-slice or a projection for 3D z-stacks data, and/or crop data to reduce memory usage; 2) automatic segmentation/tracking of multiple positions and/or multiple time-frames (batch-processing) using the two embedded neural network models YeaZ and Cellpose. YeaZ was specifically developed for yeast cells, while Cellpose is a generalist model and can segment different organisms (e.g., hematopoietic stem cells); 3) (B) Screenshot of the main GUI, where the user visualizes and corrects the result of automatic segmentation and tracking. Almost all the available functions (such as brush, eraser, edit ID or auto-separate cells) are easily accessible from a button on the top toolbar, while sliders under the left image allow quick visualization of a specific position, frame, or z-slice. To enhance visualization of the signal in the left image, the user can adjust the intensity levels with two vertical sliders on the left side of the GUI. (C) Example of the output table generated by cell cycle annotations. The annotations are saved in the broadly used CSV format allowing for quick import into GUI- or script-based spreadsheets software. The information saved includes the frame number, the cell ID, the cell cycle stage (either “G1” or “S/G2/M”), the generation number (automatically increased when division is annotated), the relative ID of the assigned parent cell, the relationship with the relative ID (either “mother” for both mother cells and cells in G1, or “bud” for buds that did not divide yet), the frame when the cell emerged and divided, and whether the history of the cell is fully known or not. Examples of cells with history not fully known are cells already present at frame 0 and cells appearing at a specific time point from outside of the field of view. Note that “is_history_known” is also visually highlighted with a question mark on the cell (e.g. cell ID 3, which was present at frame 0).

After the conversion of the image file format, the user can select any of the three following steps (all written in Python): a) a GUI for multiple data preparation steps (aligning frames, cropping images, determining background area, and selecting z-slice or projection for the segmentation step), b) automated segmentation and tracking using state-of-the-art neural networks, and c) a GUI for semi-automated correction of segmentation and tracking errors, plus annotations of budding yeast cell cycle and pedigrees (Figure 1B and Movie S1).

While it is possible to perform manual segmentation from scratch in the GUI, we highly recommend using the automated segmentation and tracking script. We embedded two of the most accurate neural networks that were recently published: YeaZ for yeast cells, and Cellpose for multiple model organisms (bright-field and phase contrast). The modularity of the code will allow for easy and quick implementation of any other segmentation algorithm (traditional or deep-learning-based) in the future.

Alongside segmentation and tracking functionalities, the GUI has an additional working mode: pedigree and cell cycle annotations. This is a functionality specifically developed for the cell cycle analysis of budding yeast cells, but it could be easily adapted to other model organisms. Annotations of the yeast cell cycle include two main steps: a) assigning the bud to the correct mother cell, and b) annotating the cell division event. Annotations are stored in a tabular format (Figure 1C) that allows reconstruction of the entire pedigree of each single cell and downstream extraction of data of interest.

Independent of whether the user decides to use the cell cycle annotation functionality, Cell-ACDC produces comprehensive output data on a single-cell level. The extraction of metadata from raw files mentioned above allows for the approximation of volumes based on the segmentation masks. Analysis of additional (fluorescence) channels enables the calculation of several quantities, such as amount, concentration, or median signal strength of the fluorescent markers. Finally, the annotation of the cell cycle additionally allows analysis of those quantities in context of the cell cycle and calculation of time-dependent properties such as growth rates.

### Overview of functionalities

The recent advancement in deep-learning-based segmentation algorithms greatly reduced the segmentation error rate, but unfortunately many times it is still required to visually inspect and correct these errors. This is a tedious and time-consuming process, especially for live-cell imaging experiments where an error in one frame requires correction of all the future frames (often hundreds of frames). For this specific step, the GUI needed to be fast, intuitive, responsive, and interactive. To allow easy detection of potential errors, we included visual help directly displayed on the images and segmentation masks while navigating through the frames (Figure 2A and Movie S2) including cell contours, cell ID, cell cycle information, lost and new cell’s contours. We automated propagation of manual corrections to future and past frames along with continuous tracking of the segmented cells while maintaining consistency with already annotated parts of the data. We implemented automated and semi-automated functions to allow quick and accurate correction once an error is detected. While we here only highlight a few examples, we explain each function in detail in the manual (Supplementary information).

**Fig 2.**
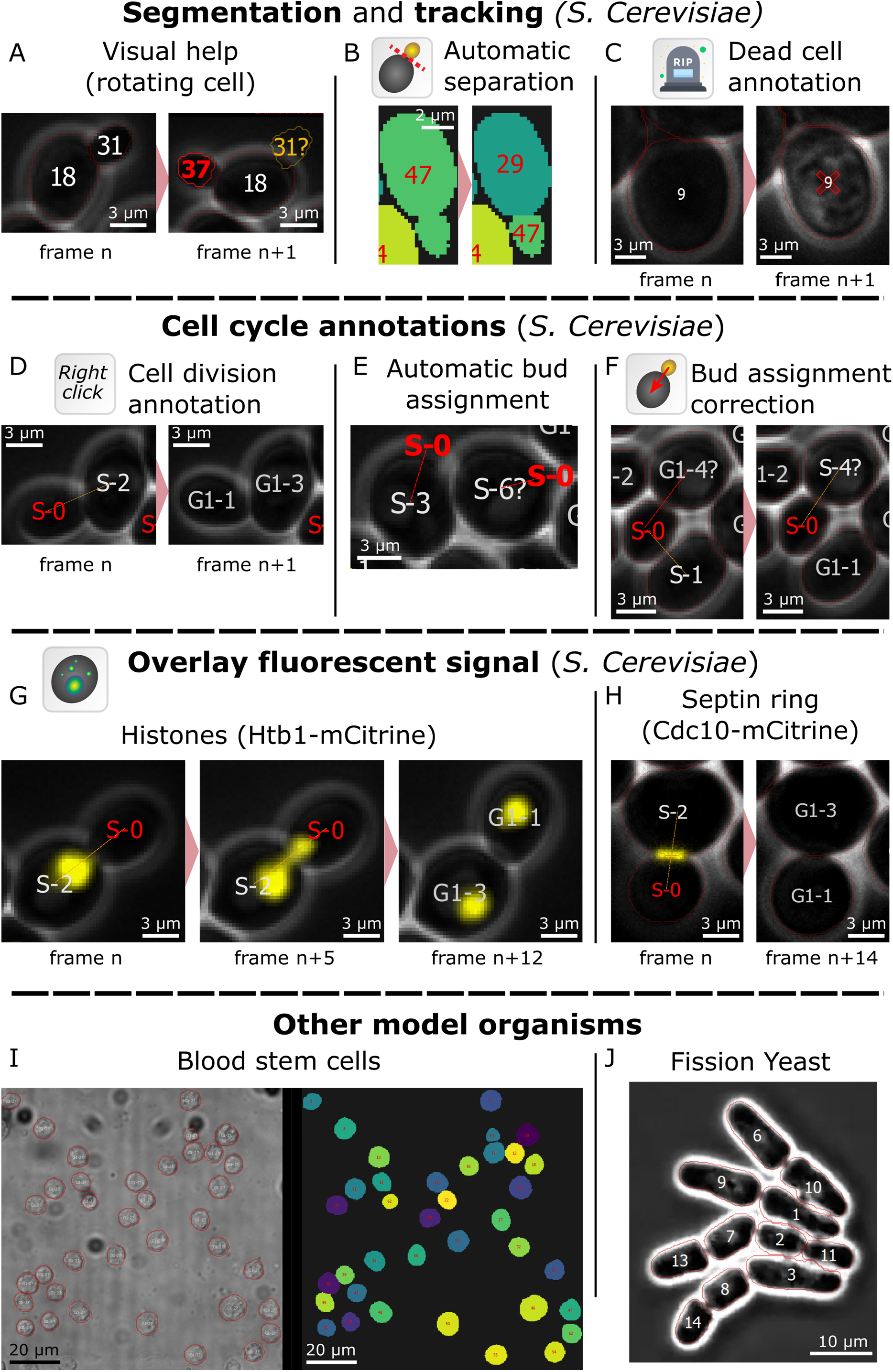
Examples of Cell-ACDC functions. (A) Visual help: information such as the cell ID, the cell cycle stage, and the generation number, as well as the segmentation contour are conveniently displayed on the cell image. Information is colour-coded: red for newly emerged/appeared cells, white for cells already present in the previous frame, and yellow for disappeared cells. This allows for quick identification of tracking errors since often lost cells are caused by an ID misplaced due to the tracking algorithm failing. (B) Automatic separation: With a single click on the merged cells, the user can trigger automatic separation. With a combination of convexity defects detection and contour approximation, the algorithm separates the cells along the predicted plane. (C) Annotate cell as “dead”: A cell can be annotated as dead with a single click, and it is then considered dead for all future frames. The user can always annotate the cell as not dead at any point in future frames. (D) Annotate cell division: Cell division is often visible due to a sudden movement of the bud. The user can then click on the cell that divided to annotate it. The related information, such as generation number and cell cycle stage, is then automatically updated for both the mother and daughter cell. This annotation can be undone at any time point in past or future frames and all the information in all the involved frames is automatically updated. (E) Automatic mother-bud pairing: When a new cell appears, an automatic assignment algorithm is triggered. Using a cost-optimization routine, the new cell is assigned to the predicted mother. (F) Mother-bud pairing correction: When the automatic mother-bud pairing fails, the user can correct the assignment with a drag and drop gesture. This can be done at any time-point of the life of the new cell and the pairing is automatically corrected on all the relevant past and future frames. (G) Overlay fluorescent signal from tagged histone Htb1. If available, the user can overlay a fluorescent signal. This is helpful, if, for example, the tagged gene is a cell cycle marker that can aid cell cycle annotations. (H) Overlay fluorescent signal from tagged septin ring (Cdc10). (I) Representative images of murine hematopoietic stem cells segmented based on bright-field signal using Cell-ACDC (based on Cellpose, using the median z-projection). (J) Segmentation using Cell-ACDC (based on YeaZ) of fission yeast (*S. Pombe*). Data from ^29^.

A typical time-consuming correction is editing the ID of an object when tracking fails. Since most of the tracking algorithms track objects based only on the previous frame, a tracking error at one frame results in the error being propagated through all preceding frames in the video. The Cell-ACDC GUI provides a real-time tracking mode that is activated when browsing through unseen frames. This allows for seamless correction of tracking errors while analysing the video frame-by-frame. Another typical segmentation error occurs when two cells are segmented as a single object (usually a mother cell with a small bud, Figure 2B and Movie S3). For this specific case, we developed a custom algorithm for automatic separation of the merged cells. With a combination of convexity defects detection and contour approximation, the cells are automatically separated. We compared this method to classic distance transform followed by watershed separation that is implemented in YeaZ, and we found consistently better performance in these specific cases (Fig. S1). If automated separation fails, the user can separate the cells manually with a dedicated function. An additional fundamental requirement is the possibility to annotate images. We implemented a variety of functionalities to annotate specific cell states, such as “dead” (Figure 1C) or “excluded from the analysis” that are activated with a single click on the cell. The corrected annotation is automatically propagated to all future frames and can be undone at any time point.

After acquiring time-lapse microscopy data of proliferating budding yeast, a typical analysis involves annotating budding events, divisions events, and identifying mother-bud pairs (Movie S4). For these specific steps, we developed three main actions: a) annotation of cell division time point (Figure 2D), b) automatic bud assignment (Figure 2E), and c) semi-automated bud assignment correction (Figure 2F).

Without the use of a cell cycle marker, cell division is often visible in the phase contrast images due to a sudden movement of the bud. To annotate this event, the user simply clicks once on the mother or bud. Cell-ACDC will automatically update the annotations table by changing the cell cycle stage and increasing the generation number to keep track of how many times a cell budded. Many times, this event is clearly visible, but other times it requires careful inspection to spot a subtle movement indicating cell division. To spot the event in this case, the user must constantly jump back and forth between frames. Therefore, responsiveness and speed of displaying data are fundamental. To achieve this, we used the high-performance python library *PyQtGraph* for the GUI elements. Furthermore, in practice it is often necessary to correct a cell division annotation multiple times. Therefore, Cell-ACDC automatically propagates corrections to all involved frames.

An important objective of cell cycle analysis with budding yeast is assignment of newly emerging buds to the correct mother cell. Using a cost optimization routine Cell-ACDC automatically assigns each emerging bud to the predicted mother cell. For all new buds, the algorithm calculates the cost of assigning the bud to any cell in G1 (i.e., cells that are not budding now). If the minimum Euclidean distance between contour pixels of the bud and a cell in G1 is low, then the cost of assigning to that cell is low and vice versa. Minimizing the cost of assignment results in automatic mother-bud pairing. To quantitatively benchmark mother-bud pairing accuracy (percentage of correctly assigned buds), we tested the algorithm with time-lapse data in three different scenarios: a) data automatically segmented with YeaZ without any correction of segmentation and tracking errors, where all the cells are eligible mothers; b) data with correction of segmentation and tracking errors, where all the cells are eligible mothers; c) data with correction, where only cells in G1 are eligible mother cells. Note that c) is the scenario in which the algorithm is currently used. In scenario a) and b), we obtained an accuracy of 67.5% and 75.5% (n=120) respectively. In scenario c), we obtained an accuracy of 90.5% (n=147). In some cases, the assignment fails, e.g., when a bud emerges close to another cell in G1 that is not the mother cell, or due to errors in earlier frames (e.g., when the correct mother cell is not annotated as being in G1). In this case, the user is notified of the potential loss of data integrity and can visit past frames to correct relevant errors. Moreover, it is sometimes not possible to determine the correctness of the assignment on the current frame, and the correct pairing is visible only after the bud has increased its size. Manually correcting such assignments would require correcting many frames where the bud must be assigned to another cell in G1 and reverting the wrong mother’s cell cycle stage back to G1. Again, the automated correction propagation is a key feature that facilitates rapid annotation.

Additionally, while it is possible to annotate the cell cycle stage using only phase contrast signal, this step can be facilitated by a fluorescent marker, such as tagged histone (e.g., Htb1 in yeast, Figure 2G) to follow the segregation of the nucleus from the mother to the bud^27^, or the septin ring (e.g. Cdc10, Figure 2H) to determine cytokinesis events^28^. To allow visualization of such fluorescent cell cycle markers, we implemented an overlay function, turned on using a button on the toolbar.

Finally, Cell-ACDC provides a framework for segmentation/tracking of other organisms such as hematopoietic stem cells (Figure 2I) or the fission yeast *S. pombe* (Figure 2J). Testing Cell-ACDC on fission yeast, we found that also for symmetrically dividing cells, cell cycle annotations and pedigree analysis are possible: after division, the two daughter cells are automatically paired, and through tracking linked to their predicted mother cell. The user can use the already implemented features to annotate division and correct daughter pairing.

### Image analysis of 3D z-stacks

Cell-ACDC offers full support for the segmentation and analysis of 3D z-stacks. Furthermore, together with the neural network Cellpose, it is possible to segment cells of various model organisms. To validate the entire image analysis pipeline including the use of 3D z-stacks, we first analyzed single time-point images of budding yeast. Using a strain expressing the fluorescent protein mKate2 from an *ACT1* promoter, we imaged both phase contrast and mKate2 signal (Fig. 3A). Secondly, using the Data Prep GUI (automatically called when segmenting 3D z-stacks), we visually selected the optimal z-slices or the projection mode. Thirdly, we segmented cells (using batch processing capabilities of Cell-ACDC) in the phase-contrast signal using the neural network YeaZ, and the mKate2 signal using Cellpose. Finally, we calculated the cell volume (see Methods section) for both segmentations. We found a strong correlation between the cell volumes calculated with the two methods (Figure 3C, Pearson’s correlation=0.98, p-value<10^−10^), indicating a strong match between cell volume estimates obtained from two different channels using two different neural networks.

**Fig 3.**
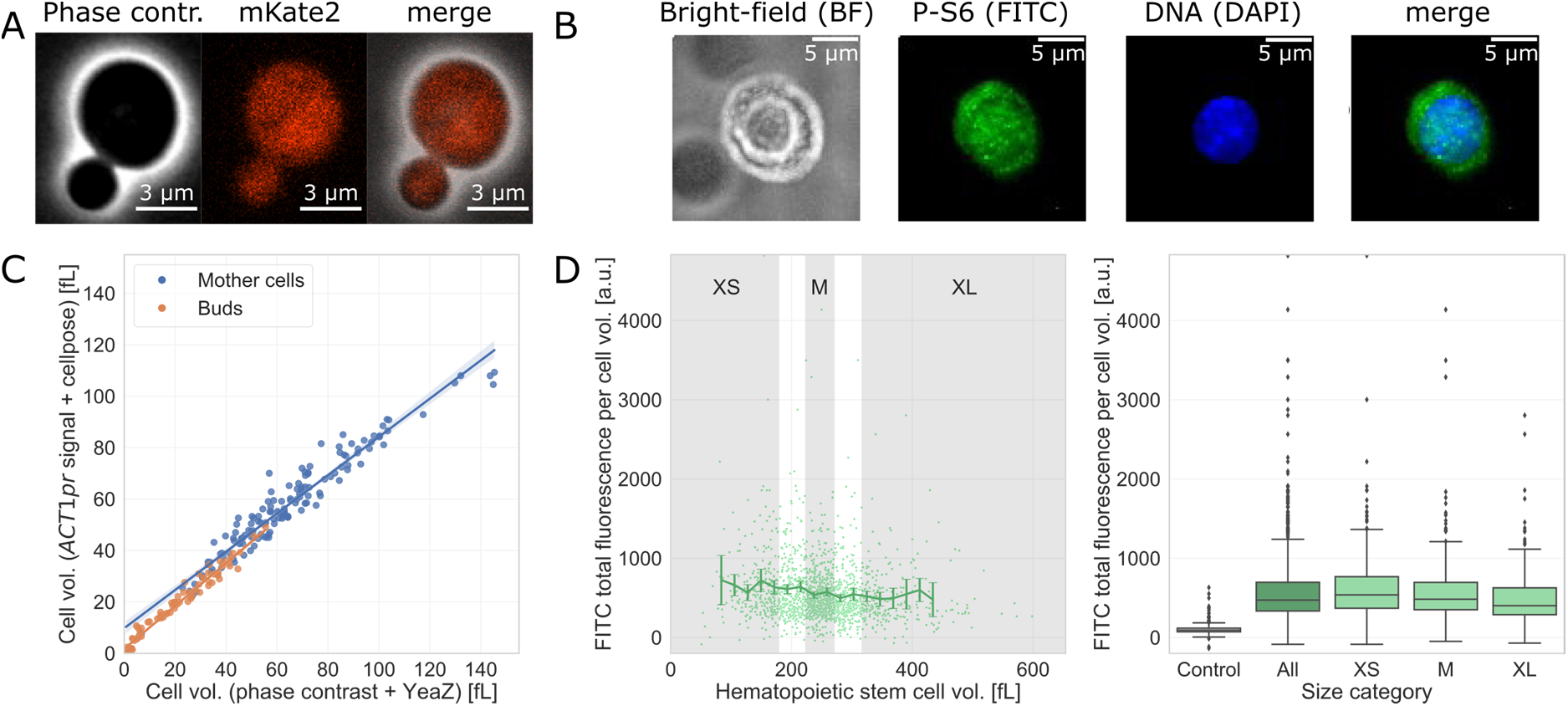
Results from analysing 3D z-stacks. (A) Representative images of the strain carrying *ACT1pr-*mKate2. (B) Representative image of hematopoietic stem cells from wild-type mice stained for P-S6 (FITC) and DNA (DAPI, scale bar is 5 μm). (C) Correlation between budding yeast cell volume calculated from cells segmented with two different methods. The cell volume is calculated from 2D segmentation masks (see ^31^), which were obtained with two different methods: segmentation with YeaZ on phase-contrast signal and segmentation with Cellpose on the fluorescent signal of mKate2 expressed from an additional *ACT1* promoter. We observe a strong agreement between the two methods (Pearson’s coefficient=0.98, p-value<10^−10^). (D) Values obtained from Cell-ACDC segmentation (using Cellpose): FITC total fluorescence intensity per cell volume (a.u.) of phospho-S6 ribosomal protein (Ser240/244) staining as a function of the HSC volume (fL). Gates of XS-, M- and XL-sized HSCs are indicated in grey (*n* = 1626 cells). FITC total fluorescence intensity per cell volume (a.u.) of phospho-S6 ribosomal protein (Ser240/244) staining in control (no primary antibody), all HSCs, XS-sized HSCs, M-sized HSCs and XL-sized HSCs (*n* = total of 234 per condition).

To validate the capabilities of Cell-ACDC to segment other model organisms and extract automatically calculated metrics from the fluorescent signal, we segmented hematopoietic stem cells (HSCs) from an immunofluorescence staining for phospho-S6 ribosomal protein (Ser240/244) using bright-field images to determine cell volume (Fig. 3B). Based on this segmentation, we evaluated the total FITC fluorescence intensity divided by cell volume, which is a readout for mTOR activity (Fig. 3D). Our results validate manual measurements showing that mTOR activity stays largely constant with increasing HSC volume^30^. Overall, these results demonstrate that Cell-ACDC enables the reliable and efficient analysis of fluorescence images of murine hematopoietic stem cells.

### Image analysis of single-live-cell imaging experiments

To validate the image analysis pipeline for live-cell imaging assays and highlight the potential for quantitative analysis of fluorescence microscopy signal, we decided to re-analyze time-course images of a yeast strain expressing the histone Htb1 endogenously tagged with a fluorescent reporter (mCitrine) that we have previously analyzed using a dedicated custom Matlab-script^15,32^. Histones are expressed in a cell cycle-dependent manner, with expression tightly coupled to DNA synthesis during S-phase^33^. After aligning the frames with the Data Prep GUI to correct for shifts during the time-lapse experiment, we segmented the videos with the batch processing segmentation script, using YeaZ on the phase-contrast signal. Next, we corrected segmentation and tracking errors, and we annotated cell cycle progression in the main GUI. Finally, we implemented a notebook in the popular open-source web application Jupyter Notebook^34^ to allow interactive transformation, exploration, statistical analysis and visualization of the Cell-ACDC output data. As expected^32^, by plotting the Htb1-mCitrine amount over entire cell cycles aligned at bud emergence, we observe a strong cell cycle dependence of Htb1 expression and a 2-fold increase around DNA replication (Figure 4A, n=109). Cell cycle annotations also allow comparing results at different cell cycle stages. We show that the amount of Htb1-mCitrine in single mother-bud pairs (before division) is about double the amount in single cells at birth (start of G1 phase). Moreover, confirming our previous analysis, we find that the amount of histones at a given cell cycle stage is largely independent of cell volume^32^ (Figure 4B). Going beyond our previous analysis, Cell-ACDC allowed us to separately analyze new-born daughter cells during their first cell cycle as well as older mother cells. By additional analysis of an untagged control strain, we ensured that autofluorescence is negligible compared to the histone-specific fluorescence signal.

**Fig 4.**
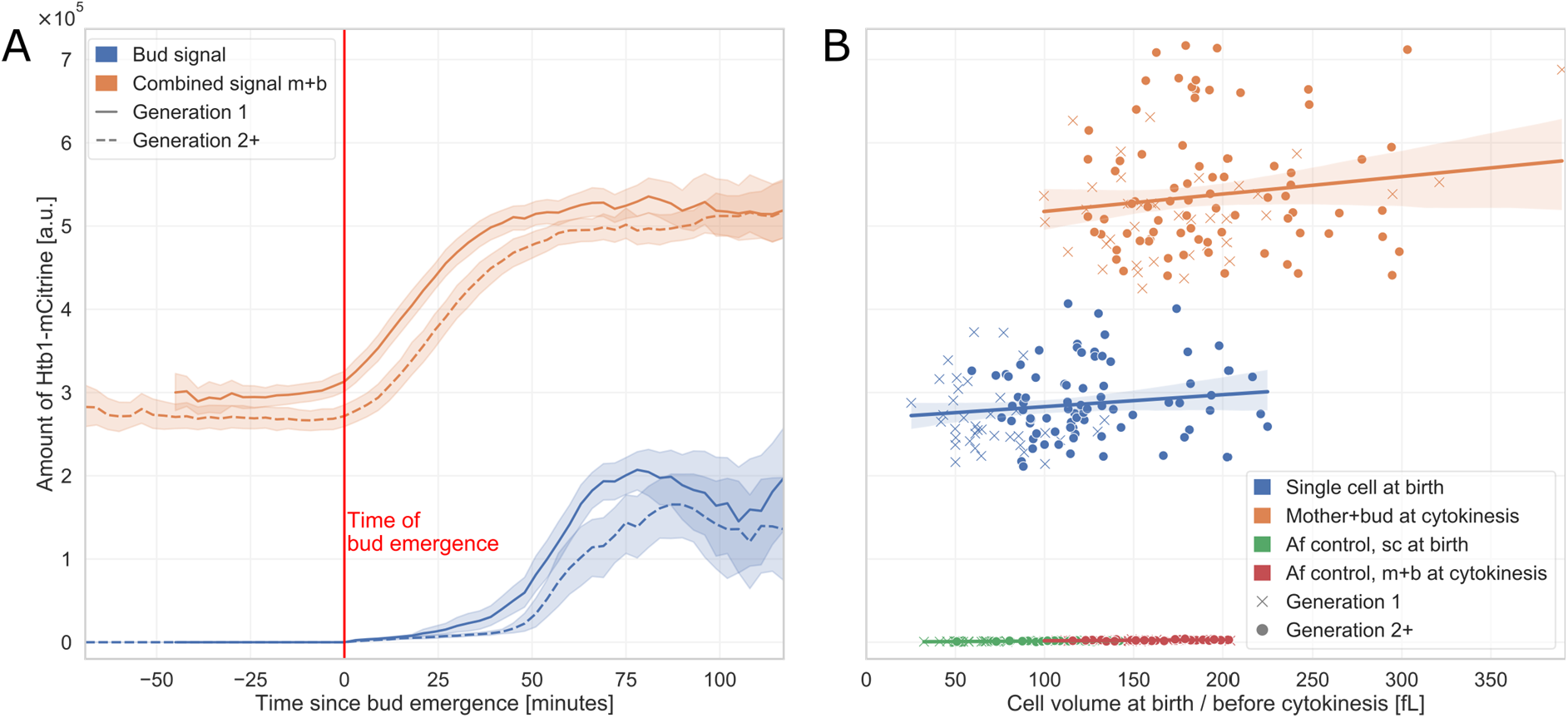
Quantitative analysis of Htb1-mCitrine expression as a function of cell cycle and cell volume. (A) Total amount of Htb1-mCitrine (total cellular fluorescence intensity after background subtraction) as a function of time for the first cell cycle of new-born daughter cells (n=38) and older cells (n=79). Single cell traces are aligned at bud emergence (time = 0). (B) Amount of Htb1-mCitrine as a function of cell volume at birth (blue) and directly before cytokinesis (combined signal of mother and bud, orange). Signal from untagged strains used as autofluorescence (Af) control is negligible. Results from (A) and (B) are consistent with our previous analysis^32^.

## Discussion

Analysis of (live)-cell imaging data is a complex task that involves several steps, some of which are often laborious and time-consuming. Despite great advances in image analysis algorithms, such as convolutional neural networks, extracting useful biological information from microscopy images can require the implementation of sophisticated pipelines. Here, we presented Cell-ACDC, an open-source, GUI-based framework that enables fast, accurate and intuitive analysis of microscopy images. We provide tools for each step of the pipeline, from the raw microscopy file to the visualization of the results (Figure 1). The software is written in Python, which is freely available for all users. We embedded recent neural network models for object detection and image segmentation, YeaZ and Cellpose. While YeaZ was specifically developed for the segmentation of yeast cells, Cellpose is generalist, enabling the segmentation of multiple model organisms.

Cell-ACDC can analyse images with different dimensionalities, from a single 2D image to 3D (z-stacks or 2D images over time) and 4D images (3D z-stacks over time). We developed workflows tuned to each specific image type.

For time-stacks, we embedded a set of tools for single-cell tracking and annotation of the yeast cell cycle. Despite the great accuracy of the embedded segmentation models, it is often required to visually inspect and correct segmentation and tracking errors. Cell-ACDC was developed to enable fast and intuitive correction of these errors, with automatic handling of correction propagation to past and future frames (Figure 2). For budding yeast live-cell imaging assays, we implemented a workflow to enable annotation of the cell cycle, either from phase-contrast signal or from a fluorescent cell cycle marker (e.g., Htb1, Cdc10 etc). With a combination of automatic mother-bud pairing and semi-automatic cell division annotation, Cell-ACDC enables accurate and fast annotation of the cell cycle stage for pedigree analysis.

For z-stacks, the user can select a specific z-slice or a projection to use for segmentation. Converting a z-stack into a 2D image is required for the neural network models. We calculated the cell volume of 228 single yeast cells using two methods: a) segmentation of a specific z-slice of the phase contrast signal using YeaZ, and b) a mean z-projection of a fluorescent marker (mKate2 expressed from an additional *ACT1* promoter) using Cellpose. We found a strong correlation between the volume calculated with the two methods (Figure 3), showing the flexibility of the segmentation pipeline.

To highlight the possibility to analyse multiple model organisms, we applied the Cell-ACDC analysis pipeline to stem cells. Our results demonstrate that Cell-ACDC provides a tool for the unbiased and efficient analysis of fluorescent images of hematopoietic stem cells, which is easy to use for researchers that do not have experience in using python. Importantly, Cell-ACDC allows for automated image analysis, which can still be manually adjusted, providing a significant advantage over manual analysis by taking less time from the researcher. In summary, Cell-ADCD is a novel and used-friendly program for the analysis of cells as diverse as budding yeast and murine hematopoietic stem cells.

### Future developments

We developed Cell-ACDC with a community-centred approach, by implementing suggestions from other research labs. We will keep this approach and when adopted by a larger community we envision a tool that can standardize live-cell imaging data processing and handling, greatly facilitating data sharing between different labs. Thanks to its modular backend, Cell-ACDC allows easy and fast implementation of image analysis models that will be developed in the future. Additionally, we plan to expand Cell-ACDC capabilities to support multiple channel segmentation and annotation of additional events (e.g., nuclear division). We also plan to implement full support for symmetrically dividing cells, with a complementary automatic sister-pairing algorithm and division annotation.

Finally, since image segmentation is often the first step in the image analysis pipeline, standardizing it will enable the development of more sophisticated downstream analysis methods (e.g., for sub-cellular feature extraction) that will be directly compatible with the output data generated by Cell-ACDC.

## Materials and methods

### Software language and packages

The software is written in Python, freely available to all users. The code is open-source, and it is available at the GitHub repository https://github.com/SchmollerLab/Cell_ACDC. For automatic conversion of raw microscopy files into the required data structure, we embedded Java Runtime Environment (automatically downloaded) and *python-bioformats*^13^ to run the popular Bio-Formats^26^ library directly from Python. Thanks to a GUI-based wizard, the user can automatically generate the required data structure. The GUI backend is written using PyQt, a set of Python bindings for the Qt cross-platform C++ framework. Qt is a platform specifically designed for the development of fast and intuitive GUIs. To ensure a smooth user experience the images and the annotations are displayed using PyQtGraph, a Python library specifically designed for interactive and fast displaying of data. To ensure easy installation of Cell-ACDC, we provide ready to use virtual environments with the two most popular package installers, Anaconda and Pip. Finally, we provide a Quick Start Guide to start using Cell-ACDC as fast as possible and a User Manual that extensively documents every single function available.

### Live cell imaging

Fig. 2C shows strain KSY306-3 (*Mat a, his3::LexA-ER-AD-TF-HIS3 whi5::kanMX6-LexApr-WHI5-ADH1term-LEU2 exo84::EXO84-mCirine-ADH1term-cglaTRP1 cdc10::CDC10-mNeptune2*.*5-ADH1term-ADE2*) growing on SC media with 2% glycerol and 1% ethanol (SCGE) after pre-culture in SCGE with 20 nM β-estradiol. Data displayed in Fig 2D-H and 3A was obtained from raw microscopy files included in our previous publication^31^. Specifically, Fig. 2C-H shows the strain KSY234-1; Fig. 3A shows strain KSY282-2. Briefly, live-cell time-lapse microscopy was performed using a Nikon Eclipse microscope equipped with a plan-apo λ 100×/1.45NA Ph3 oil immersion objective. Cells were imaged in a custom made microfluidic device made of polydimethylsiloxane and a glass coverslip. A flow of 40 μl/min of synthetic complete liquid medium with 2% glucose was constantly applied at 30°C.

Data displayed in Fig 1, Fig 2A-B-D-E-F-G and Fig 4 was obtained from raw microscopy files included in our previous publication^32^. The diploid strain KCY050-2 carries endogenously tagged Htb1, while strain ASY020-1 was used as autofluorescence control^32^. Live-cell time-lapse microscopy was performed using a Zeiss LSM 800 microscope equipped with a plan-apochromat 40x/1.3NA oil immersion objective coupled to an Axiocam 506 camera. Note that Fig4B here in essence reproduces Fig 1c of publication^32^. However, a different subset of cells from the raw data was analysed.

### Fluorescence staining of hematopoietic stem cells

Murine bone marrow (BM)-derived live G0/1 HSCs (Lin^-^, Sca1/Ly6^+^, CD117/cKit^+^, CD150/Slamf1^+^, CD48/Slamf2^-^, 7-ADD^-^) were isolated as described previously^30^. Briefly, BM was harvested by flushing the long bones. Red blood cells were lysed in ammonium-chloride-potassium (ACK) buffer and samples were washed in Iscove’s Modified Dulbecco’s Medium (IMDM) containing 2 % fetal bovine serum (FBS). BM cells were resuspended at 10^6^ cells/mL in pre-warmed IMEM supplemented with 2 % FBS and 6.6 μg/mL Hoechst-33342 (Thermo Fisher Scientific, #H3570). After 45 min of incubation at 37°C in a water-bath, cells were washed with cold IMEM with 2 % FBS and kept at 4°C. Lineage positive cells were depleted using a mouse lineage cell depletion kit and the following antibodies were used for staining: Rat monoclonal PE anti-mouse CD150, BD Biosciences, Cat#562651; RRID: AB_2737705; Rat monoclonal APC anti-mouse CD117, BD Biosciences, Cat#561074; RRID: AB_10563203, Armenian hamster monoclonal APC/Cy7 anti-mouse CD48, BioLegend, Cat#103431; RRID: AB_2561462, Rat monoclonal BV711 anti-mouse Ly-6A/E, BioLegend, Cat#108131; RRID: AB_2562241. Cells were sorted using an Aria cell sorter (Becton Dickinson).

For immunofluorescence analyses, Fisherbrand™ Superfrost™ Plus Microscope Slides were primed with 0.1 % polylysine for 5 min, washed with dH_2_O and air-dried. HSCs were distributed on slides and incubated for 1 h in a humidified chamber at RT. HSCs were fixed for 20 min at RT with freshly prepared 4% paraformaldehyde (PFA, pH 7.2) and then washed three times with PBS. HSCs were permeabilized for 20 min in 0.2 % Triton-X 100, washed three times with PBS, and blocked for 30 min using 10 % Donkey Serum (Sigma) in PBS. Cells were incubated with primary antibody in 10 % Donkey Serum in PBS overnight at 4 °C: Phospho-S6 Ribosomal Protein (Ser240/244) Rabbit mAb (Cell Signaling Technology, Cat# 5364; RRID:AB_10694233). After HSCs were washed three times with PBS + 0.1 % Tween-20, the secondary antibody solution (1:500, goat anti-rabbit Alexa 488, Cell Signaling Technology, 4412S) was added for 1 h at RT in the dark in 10 % Donkey Serum in PBS. Coverslips were mounted with ProLong Gold Antifade Reagent with (Invitrogen, Molecular Probes) and imaged after 12 h. Control slides were not treated with primary antibody. Images were acquired using a DeltaVision Elite microscope (Applied Precision) platform (GE Healthcare Bio-Sciences) equipped with a CoolSNAP HQ2 camera (Roper), 250W Xenon lamps, SoftWoRx software (Applied Precision). Deconvolution was performed using SoftWoRx software with default settings. Cells that were 2.5 times larger than the mean were excluded from the analysis. To analyse HSCs of a specific size, the evaluated the 10 % smallest (XS-HSCs), the 10 % largest (XL-HSCs) and +/- 10 % HSCs of mean size (M-HSCs).

### Cell volume calculation

Cell volume is estimated from 2D segmentation masks as follows: a) the object is aligned along its major axis, b) the volume of each horizontal slice with 1 pixel height is calculated assuming rotational symmetry along the slice’s middle axis, c) volumes of the slices are added to obtain the final volume. We previously reported^31^ that for budding yeast, this method well agrees with alternative methods, such as 3D reconstruction from z-stacks using confocal microscopy.

### Downstream analysis

For downstream analysis, we provide a notebook, written in python in the popular data science tool Jupyter Notebooks^34^. The user can select files to analyse by following a series of prompts and file dialogs, which also enables data pooling and comparison of subsets such as different strains or different conditions. The files selected are then iteratively loaded and geometric properties (e.g., area, solidity, elongation) are calculated using the package scikit-image^35^. Those quantities are complemented by additional parameters specific to time-lapse experiments, including cell age at frame *n*, growth rate, G1 and S/G2/M durations, as well as the first and last frames of cell appearance. In addition, signal amount and concentration for all available fluorescence channels are calculated. For this, the mean signal is corrected for background fluorescence by subtracting the median signal of all background pixels, which are determined as non-cell areas based on the cell segmentation masks. We define signal amount as corrected mean multiplied by the area of the segmented object (in pixels) and the signal concentration is obtained by dividing the amount by cell volume (calculated as described above).

We then perform two data aggregations using functions of the package pandas^36^. First, we connect the mother cell data with data of the respective buds and obtain single-cell time traces using the cell IDs. Second, we use generation number and cell cycle stage information to calculate cell-cycle-specific data.

Figure 3 was created using the output from ACDC without any pre-processing. Cell volumes and FITC concentrations were calculated as described above. In Figure 4A, all individual cell cycle traces are aligned at bud emergence. To obtain the combined signal of mother cells and their buds, we summed the respective fluorescence signal amounts

### Continuous tracking

For the continuous tracking of single cells in the main GUI, we developed a cost-optimization routine to determine the optimal assignment of the segmented objects between two consecutive frames. Firstly, a cost matrix **C** is computed: given a list **x** of object IDs [*id*_0_, *id*_1_ *… id*_*n*1_] in frame *n* − 1, and a list **y** of [*id*_0_, *id*_1_ *… id*_*n*2_] in frame *n*, each element *c*_*i,j*_ is equal to the intersection over area (*IoA*) score between the *i*^*th*^ object of list **y** (*id*_*yi*_) and the *j*^*th*^ object of list **x** (*id*_*xj*_). The *IoA* is calculated as the number of intersecting pixels between *id*_*yi*_ and *id*_*xj*_ divided by the area of *id*_*xj*_. Next, any object with maximal *IoA* score less than 0.4 is considered a new object (e.g., a newly emerging bud), and receives a new *id*_*new*_ ∉ **x** ∧ *id*_*new*_ ∉ **y**. The remaining objects from frame *n* are assigned as follows: each unassigned object of list **y** is assigned to the object of list **x** with maximum *IoA* score unless the object from list **x** has a higher *IoA* with another object from list **y**. After having assigned objects from frame *n* to all objects from frame *n* − 1, the remaining objects are considered new and receive a new *id*_*new*_. Since the tracking algorithm is embedded into the main GUI, the key aspect is the execution speed (e.g., to spot subtle movements of a bud that indicate a cell division event). Therefore, we benchmarked it with a segmentation mask containing 99 cells to be tracked, and we calculated the average execution speed after 1000 runs. Our algorithm, on average, took about 45 ms, while the tracking algorithm embedded in the YeaZ model took about 260 ms. This is a considerable improvement that enhances the overall speed when navigating through frames in the main GUI.

### Automatic separation of merged objects

Another algorithm embedded into Cell-ACDC is the automatic separation of merged objects. Since both Cellpose and YeaZ provide methods for separation, we developed our algorithm to provide an additional option for cases where Cellpose or YeaZ failed. The goal of the method is to separate the object along a restriction site. Firstly, the contour of the object is approximated to avoid spurious separation planes due to irregularities in the contour shape line. This is achieved with the OpenCV (computer-vision library for Python)^37^ function `approxPolyDP` using 10% of the contour length as the epsilon parameter. Next, the convexity defects of the convex hull of the approximated contour are computed using the OpenCV function `convexityDefects`. Finally, if the detected defects are two, then the object is separate along the line connecting the two convexity defects.

### Automatic mother-bud pairing

When the GUI is in “cell cycle analysis” mode, every new object appearing in the next frame is considered as a bud that needs to be assigned to a cell in G1 (if not already assigned in a previous visit of the frame). Firstly, the algorithm determines if there are enough cells in G1 for the new cells, and if not, a warning is triggered and the user can decide to automatically annotate that the history of these cells is not known (e.g., a cell appearing from outside of the field of view), or can annotate previous divisions of cells to increase the number of cells in G1 (if, for example, a division event was missed). After this checkpoint, the contour of each cell in frame *n* and frame *n* + 1 is computed. Then, given the lists **a, b** of the new cells and old cells in G1, respectively, a cost matrix **C** is calculated. Each *c*_*i,j*_ element is equal to the minimum Euclidean distance between the pixels of the *a*_*i*_ cell’s contour and the pixels of the *b*_*j*_ cell’s contour. The optimal assignment is calculated using the minimum weight matching in bipartite graphs routine called `linear_sum_assignment` implemented in the Python package SciPy^38^.

### Automatic propagation of corrections to future and past frames

One of the most tedious and time-consuming processes is the correction of the same error when it appears in many consecutive frames. To speed-up this process we developed a series of routines to automatically propagate the correction to all the relevant future and past frames, when possible. Automatic propagation is triggered in the following situations: a) mother-bud pairing correction, b) cell division annotation and its correction, c) tracking error correction, d) object deletion, e) editing a cell’s ID, f) excluding a cell from analysis, and g) annotating a dead cell. For the situations c)-g), the user can choose between applying the same correction/annotation to all future frames or simply repeat tracking for all the future frames. For the situations a) and b), the propagation is completely automatic. The correction of mother-bud pairing involves three cells: the bud, the wrong mother cell, and the correct mother cell. First, the correct mother cell must be a cell in G1, since the assumption is that each mother cell can have only one bud assigned to it. Furthermore, the correct mother must not have had any other bud assigned to it for all the frames in which the bud to be corrected is present. If the correct mother cell satisfies the eligibility criteria, once the user corrects the pairing, all the frames in which the annotation is wrong, are automatically corrected: the wrong mother cell goes back to the state it had before the bud was assigned to it, while the correct mother is assigned to the bud. Since correction is automatic to both past and future frames, it can be performed at any time point.

The correction of cell division annotation can be done on both a cell in G1 or a cell in S/G2/M. If the user clicks on a cell in S/G2/M (annotating division) at frame *n*, the automatic routine will annotate the division event at frame *n* for both mother and bud. Then, it will check if there are future frames that were previously annotated as cell in S/G2/M and will correct them accordingly. Otherwise, if the user clicks on a cell in G1 (undoing division annotation), the routine sets both the cell and the bud it had in the previous cycle back to S/G2/M for all the future (until the cell is in S/G2/M again or we reach the last visited frame) and past frames (until the mother cell is in S/G2/M again). Automatic propagation allows for annotating or undoing the annotation at any time point, which is particularly useful when toggling back and forth between frames is required for accurate cell division annotation.

## Acknowledgments

We thank Jacob Kim, Jordan Xiao, Christian Everett Wright, Yagya Chadha, Dimitra Chatzitheodoridou, Igor Kukhtevich, María Rocha Acevedo, Marlet Morales Franco, and Soham Bharadwaj for testing Cell-ACDC and providing valuable feedback. We thank Igor Kukthevich for imaging KSY306-3, and Fred Chang, Benjamin D. Knapp and Kerwyn Casey Huang for providing fission yeast data. This work was supported by the DFG through project SCHM3031/6-1, by the Human Frontier Science Program (career development award to K.M.S.) and by the Helmholtz Association.

B.M. is supported by the Helmholtz Association under the joint research school Munich School for Data Science - MUDS.J.L. is supported by Academy of Finland.

