## Supplementary Information for "Cell-ACDC: a user-friendly toolset embedding state-of-the-art neural networks for segmentation, tracking and cell cycle annotations of live-cell imaging data"

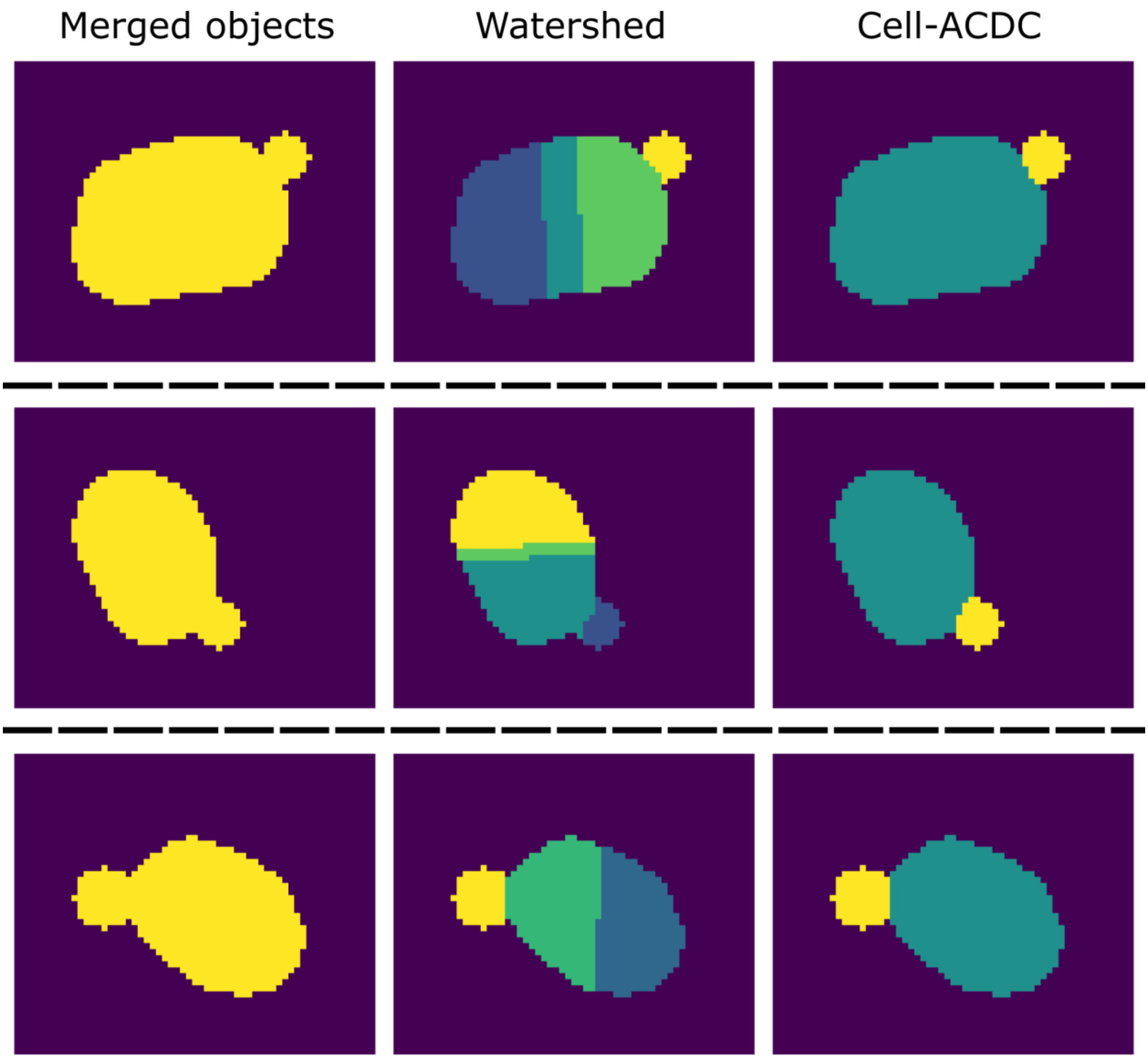


**Fig. S1 – Comparison between Cell-ACDC automatic separation algorithm and classic distance transform plus watershed.** Three example cells, for which the separation method used by YeaZ fails. Note that the YeaZ splitting algorithm resulted in the objects not being separated, but for visualization purposes we show the results of the watershed separation. The YeaZ method attempts at removing wrong watershed lines by comparing them with the prediction of the neural network model. If the predictions that the lines are part of the cell are high enough, the lines are removed, and the objects are merged again. However, in these cases, the prediction at the mother-bud neck is high as well and the objects are merged again, resulting in bud connected to the mother cell. Cell-ACDC uses a different strategy. It uses a combination of convexity defects and contour approximation to determine the constriction site and it separates along that mother-bud neck. Note that this approach works only for two objects merged and separated by a constriction.

Link: https://drive.google.com/file/d/1fLaB4irUaat3i5PKeYWEYge5pKUxfmFE/view?usp=sharing

**Movie S1 – Video of a fully annotated position with cells disappearing due to suboptimal channel width.**

Link: https://drive.google.com/file/d/1dHT6OJgl-Sfveu8RLUAwM-kgWonVMJ-h/view?usp=sharing

**Movie S2 – Visual help (rotating cell) in the main GUI**. Cell 18 rotates at frame $n + 1$ resulting in a tracking error. Thanks to the annotations on the images, the user detects that cell 31 disappears, while a new cell 37 appears. To fix this, the user can manually assign ID 31 to cell 37. If the user does not see this and tries to continue to the next frame anyway, a warning message (pop-up window) will warn the user that cell 31 was lost and he/she can decide to continue or not.

Link: https://drive.google.com/file/d/1Ya7lBrkqVg2juggZ4SJKUOW4Ucvyq7Mq/view?usp=sharing

**Movie S3 – Automatic separation of merged mother-bud.** After activating the "Automatic separation mode" with a button on the toolbar (or key shortcut), the user right-clicks on the merged objects to automatically separate them.

Link: https://drive.google.com/file/d/1fJ93wDqK020HZcUHChAXFSJMm2IVTfcv/view?usp=sharing

**Movie S4 – Cell cycle annotations example.** When annotating cell cycle information, the user must keep an eye on two events: correctness of the automatic mother-pairing and division event. In this video, the user navigates through the frames and at a specific time-point the bud with ID=4 is automatically assigned to mother with ID=1. Next, when a sudden movement of bud with ID=4 is visible, the user clicks on the mother or bud to automatically annotate the division event.
